# A phase-based mechanism of attentional selection in the auditory system

**DOI:** 10.1101/2025.05.23.655714

**Authors:** Troby Ka-Yan Lui, Victorina De Combles De Nayves, Céline Cappe, Benedikt Zoefel

## Abstract

Perceptual targets are often easier to detect and process if they occur at a predictable time. This perceptual benefit of temporal predictability suggests that neural resources are allocated to specific moments predicted to be most informative, or away from those predicted to be distracting. The sensitivity to sensory information changes with the phase of neural oscillations. An alignment of oscillatory phase according to the predicted time and relevance of expected events has therefore been proposed as a mechanism underlying attentional selection. This hypothesis predicts that opposite phases are aligned to relevant and distracting events so that the former are amplified and the latter suppressed, but such an effect remained to be demonstrated.

We presented 25 human participants with acoustic stimuli that occurred at a predictable or unpredictable time during a pitch-discrimination task. Temporal predictability did not only lead to faster reaction times, but it also aligned the phase of beta oscillations in the electroencephalogram (EEG) in anticipation of the stimulus. Crucially, the aligned phase was opposite, depending on whether the stimulus was relevant or a distractor for the task. Single-trial deviations from the mean aligned phases slowed down behavioural responses, demonstrating functional significance of the observed neural effects. All effects occurred in the absence of rhythmic stimulation, an important but rarely tested criterion to rule out spurious phase alignment from preceding events in a stimulus sequence.

We therefore conclude that the alignment of oscillatory phase underlies the selection of sensory information in time. We speculate that beta oscillations allocate neural resources in networks comprising auditory and sensorimotor regions to the expected time of auditory information, facilitating behavioural responses to upcoming events.

## Introduction

Anticipating the timing of future events is fundamental for perceptual and cognitive functions as it ensures that neural resources can be efficiently allocated to upcoming sensory information^1^. Neural oscillations reflect alternating phases of low and high neural excitability^2^ which translate into low and high sensitivity for sensory information^3^, respectively. A dominant view in several fields of research, such as those on “neural entrainment^4^” and “active sensing^5^”, is that the phase of oscillations can be adjusted according to the predicted relevance of sensory information. Accordingly, the phase of high sensory sensitivity is aligned with important events that are then amplified, and the phase of low sensitivity is aligned with anticipated distractors that are then suppressed. In such a scenario, the oscillatory phase acts as a mechanism for attentional selection^6–8^. However, experimental support for such a phase-based mechanism of attentional selection remained sparse, due to several reasons.

First, an oscillatory phase alignment to anticipated information is dominantly tested in settings with rhythmic sensory stimulation^6,9–12^. Rhythmic stimulation makes it difficult to identify a genuine alignment of phase in anticipation of an event, as the response to the preceding event in the stimulus sequence can cause spurious phase alignment^13–15^. Outside of these rhythmic contexts, Bonnefond and Jensen^16^ found that the phase of alpha oscillations (∼8-12 Hz) aligns to the predicted onsets of distractors in a visual working memory task. This phase alignment was enhanced for stronger distractors, implying that the oscillatory phase suppresses distractor processing. This effect was extended to visual perception in a later study^17^. Nevertheless, the number of studies that tested for phase alignment to temporally predictable, non-rhythmic events in the presence of attentional competition remains surprisingly low.

Second, the lack of experimental support for phase alignment to non-rhythmic events is particularly strong in the auditory domain (see **Discussion** for a more detailed overview of the literature). This is in stark contrast to the fact that audition relies more on information unfolding over time than other modalities^18,19^, and therefore requires a mechanism to select information in time. Two studies did not find auditory phase alignment to expected auditory stimuli in a paradigm that involved a competition between targets and distractors for attentional resources^17,20^. However, both temporal predictions and competition were induced cross-modally (visual vs. auditory) which might not have been sufficiently strong, or biased attention towards the often-dominant visual modality.

Third, an alignment of phase to expected events is necessary but not sufficient evidence for the described mechanism of attentional selection, as it does not show that the aligned phase is *used* to select information. Such a mechanism predicts that *opposite* phases are aligned to relevant and distracting events so that the former are amplified and the latter suppressed^6^, but such an effect remained to be demonstrated, irrespective of the modality involved.

We addressed these three points in the current study. We presented participants with non-rhythmic acoustic stimuli that occurred at predictable or unpredictable times (**Fig. 1**). These stimuli needed to be attended or suppressed to perform well in a pitch discrimination task, respectively. We found that the phase of beta oscillations in the EEG aligns to the predicted stimulus occurrence. Crucially, this phase was opposite, depending on whether the stimulus was necessary or a distractor for the task. Participants responded more slowly in trials where the neural phase deviated from its typically aligned value. We therefore report evidence for a mechanism of attentional selection in time that is based on oscillatory phase.

**Figure 1.**
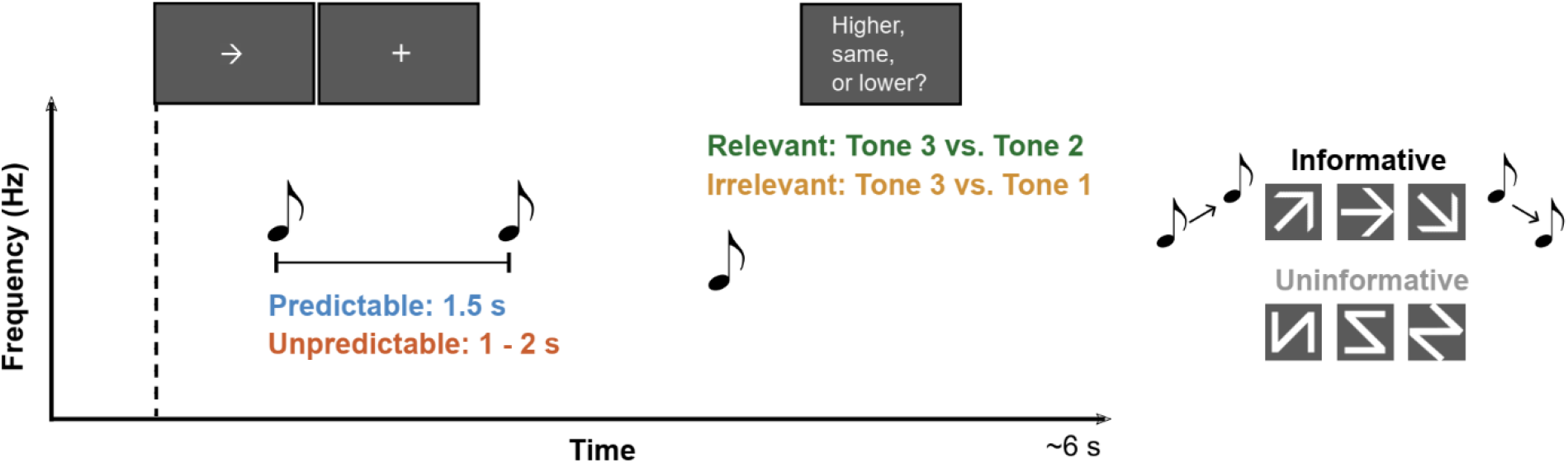
Experimental paradigm. Each trial consisted of three consecutively presented pure tones. Participants were asked to decide whether the pitch of the third tone (reference tone) was higher, lower, or identical to that of the second or first tone (comparison tone). If the second tone was the comparison tone, it was relevant for the task. If the first tone was the comparison tone, the second tone was an irrelevant distractor. The timing of the second tone was either predictable (1.5 s) or unpredictable (random between 1 and 2 s). A visual cue was presented with the first tone that was either informative or uninformative for the pitch difference between the first and second tone.

## Results

### Overview

In each trial (**Fig. 1**), participants were presented with three consecutive pure tones. They were asked to decide whether the pitch of the third tone (the “reference tone”) was higher, lower or identical to the pitch of one of the other two tones (the “comparison tone”). The second tone was therefore either necessary to do the task or an irrelevant distractor, depending on whether it was the comparison tone (“relevant condition”) or not (“irrelevant condition”). Prior to each experimental block, participants were informed about which of the tones was the comparison tone in that block. In some blocks, the delay between the first two tones was fixed to 1.5 s (“predictable condition”). In other blocks, the delay varied in each trial, selected randomly from a range between 1 and 2 s (“unpredictable condition”). We hypothesized that (1) participants leverage the predictable timing to align their phase to the anticipated onset of the second tone, (2) do so according to the tone’s relevance (opposite phases between relevant and irrelevant tones, but only if their timing is predictable), and (3) the phase alignment and opposition boosts performance in the pitch discrimination task.

We also tested whether a visual cue can help participants in their task and the associated phase alignment. Immediately before and during the first tone, an arrow showed participants whether the second tone’s pitch will be higher, lower, or identical to that of the first tone (“informative condition”), or the arrow was replaced with one out of three meaningless symbols (“uninformative condition”). We hypothesized that informative cues enhance the phase alignment to the second tone and further improve performance in the pitch discrimination task.

### Temporal predictability accelerates pitch discrimination

To improve intuitiveness of our results (higher values = faster responses), we converted reaction time to response speed (its reciprocal), but verified that none of our results depend on this decision. We used a linear mixed model to test how the predictability of the second tone’s timing, its relevance for the task, and the presence of a cue jointly determined response speed in pitch discrimination. We found that participants responded faster when the second tone was the comparison tone and therefore relevant, as compared to when it was irrelevant and the first tone was the comparison tone (t(592) = 6.58, p < 0.001, CI [0.11 0.21]; **Fig. 2A**). When the second tone was irrelevant, participants had to remember the pitch of the first tone that occurred earlier than the second tone. However, this difference in delay was controlled for by adapting pitch differences to participant’s individual thresholds, separately for relevant and irrelevant conditions (see **Methods**). We therefore conclude that participants took longer to respond when the second tone was irrelevant, because it acted as a distractor. Importantly, we also found that participants responded faster when the second tone occurred at a predictable time (t(592) = 4.79, p < 0.001, CI [0.07 0.16]; **Fig. 2A**). This result is in line with our hypothesis and suggests that participants leveraged the predictable timing to improve their discrimination performance, possibly by allocating neural resources to the expected timing of the tone.

**Figure 2.**
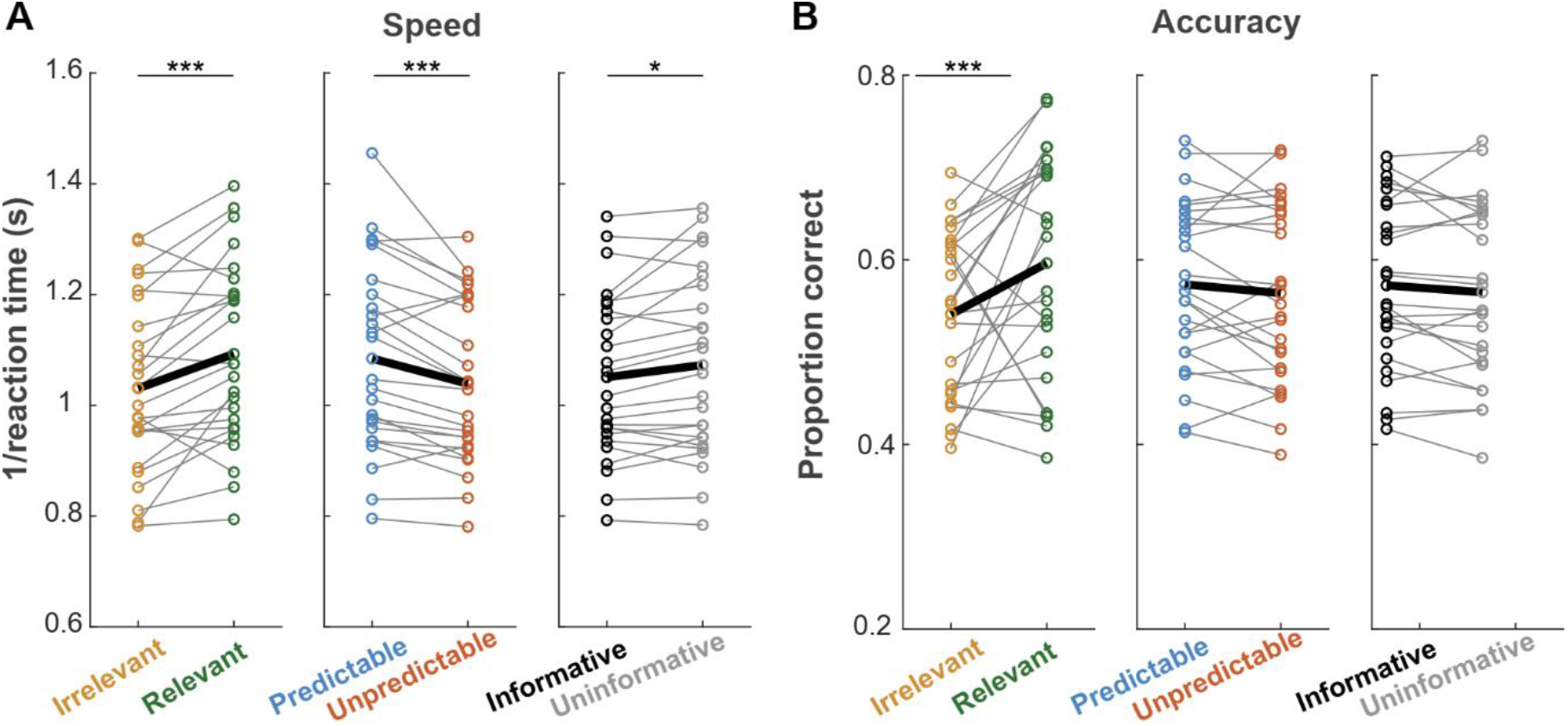
Response speed (**A**) and accuracy (**B**) in the pitch discrimination task. Circles show data from individual subjects and black thick lines illustrate their mean. *: p < 0.05; ***: p < 0.001.

Contrary to our hypothesis, participants responded faster in the presence of an uninformative cue as compared to an informative cue (t(592) = 2.32, p = 0.02, CI [0.009 0.10]; **Fig. 2A**). This could be due to the higher cognitive load when the cue is informative and needs to be processed. This cognitive load seems to outweigh the higher predictability of the second tone’s pitch during informative cues, leading to longer reaction times.

When the second tone was relevant, participants did not only respond faster, but also more accurately (t(592) = 5.63, p < 0.001, CI [0.12 0.25]; **Fig. 2B**). However, neither temporal predictability nor cue had a reliable impact on accuracy (predictability: t(592) = 0.96, p = 0.34, CI [-0.03 0.1]; cue: t(592) = 0.76, p = 0.45, CI [-0.04 0.09]; **Fig. 2B**). None of the interaction effects for reaction time and accuracy were significant (all p > 0.05).

### Temporal predictability amplifies neural responses to expected acoustic events

**Fig. 3** shows the neural responses preceding and evoked by second tone (time 0). We found that the second tone evoked a stronger response when it was temporally predictable (blue in **Fig. 3A**) than when it was not (red in **Fig. 3A;** horizontal lines show time points where a paired t-test yielded an FDR-corrected p<0.05 for channel Fz). This finding mirrors our behavioural results and suggests that a tone is represented more strongly in neural activity when its timing is predictable, and that this leads to better discriminability.

**Figure 3.**
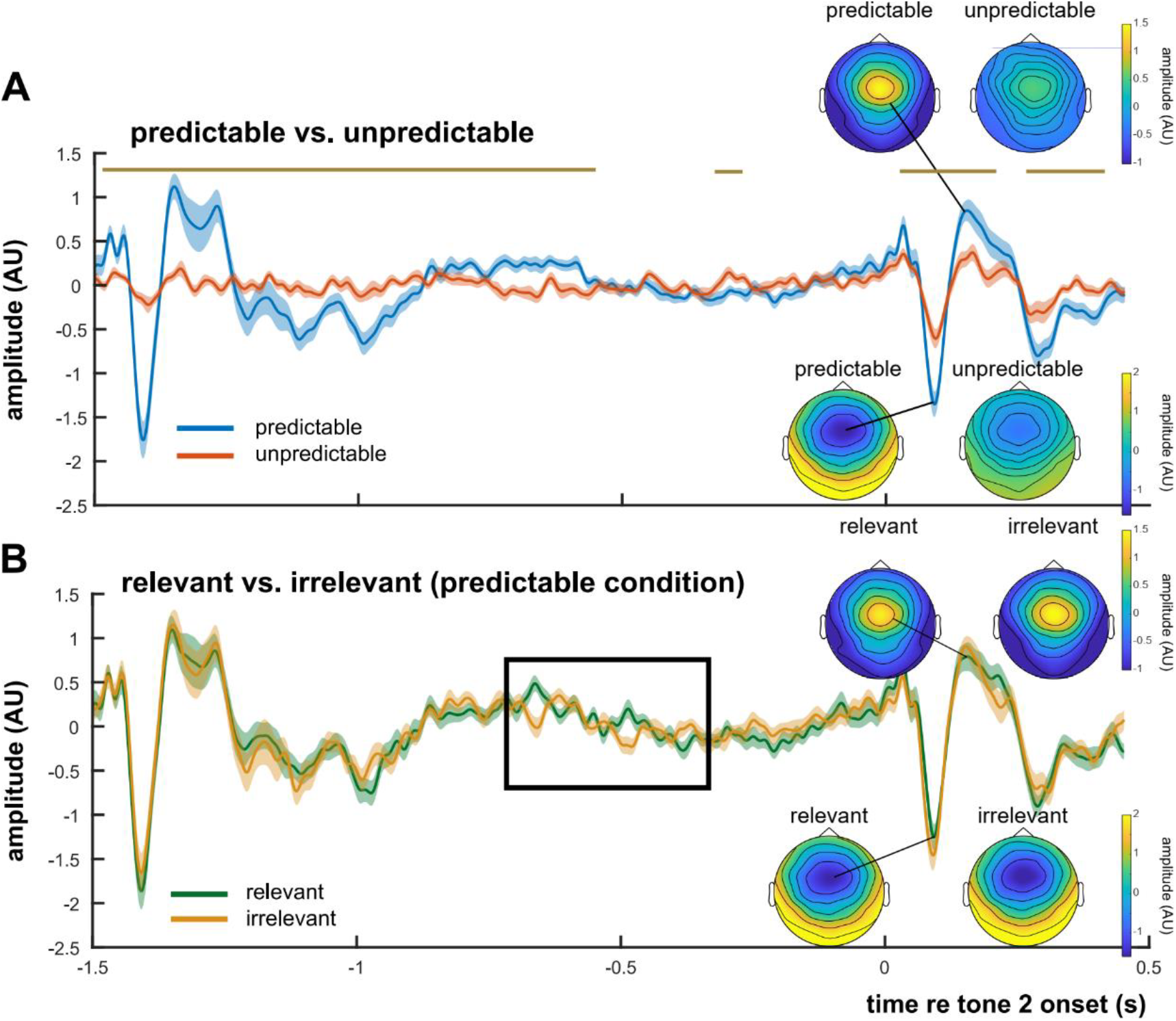
Neural response at channel Fz evoked and preceding the second tone (time 0), separately for tones with predictable and unpredictable timing (**A**), and for relevant and irrelevant (distracting) tones (**B**). Shaded areas depict the standard error of the mean (SEM). Insets show topographies for two selected time points (corresponding to N1 at +92 ms and P2 at +149 ms relative to the second tone). In **A**, the golden horizontal line shows time points with reliable difference between conditions (FDR-corrected p < 0.05). In **B**, the black box illustrates a phase opposition between conditions that is examined statistically in Fig. 4C.

In contrast to our behavioural results, relevant and irrelevant tones did not evoke different neural responses (**Fig. 3B**; all FDR-corrected p>0.05). The increased task difficulty when the second tone acted as an irrelevant distractor was therefore not directly reflected in evoked neural activity. However, we found evidence for a phase-based mechanism of distractor processing that is already visible in the EEG response preceding the second tone (black box in **Fig. 3B**), and that we will come back to below.

### Temporal predictability aligns the phase of beta oscillations to expected acoustic events

We then tested whether neural oscillations aligned their phase to the expected timing of the second tone. We quantified phase alignment using inter-trial coherence (ITC). The higher the phase consistency between trials at a given time-frequency point, the higher the ITC. **Fig. 4A** shows the results from a paired t-test contrasting ITC between predictable and unpredictable conditions. We found that ITC in the frequency range of beta oscillations increased before the tone when its onset was predictable. Averaged across EEG channels, this phase alignment was strongest at 0.5 s before tone onset and at 15 Hz (t(24) = 4.68, FDR-corrected p = 0.04). The effect was stronger for individual channels and peaked in parieto-occipital channels (e.g., PO8: t(24) = 8.17 for -0.5 s/18 Hz; FDR-corrected p < 0.001) as well as frontal ones (e.g., F1: t(24) = 5.78 for -0.51 s/17 Hz; FDR-corrected p = 0.04). The topography of this phase alignment (inset in **Fig. 4A**) resembles oscillatory phase effects in audition reported previously^21^. Explorative source localisation of the ITC contrast (−0.5 Hz/18 Hz) between predictable and unpredictable conditions (**Fig. 4B**) suggests origins that include left inferior frontal cortex and right sensorimotor cortex, both regions implicated in auditory temporal predictions^22,23^. Together, these findings illustrate that beta oscillations align their phase to temporally predictable acoustic events even if they are non-rhythmic.

**Figure 4.**
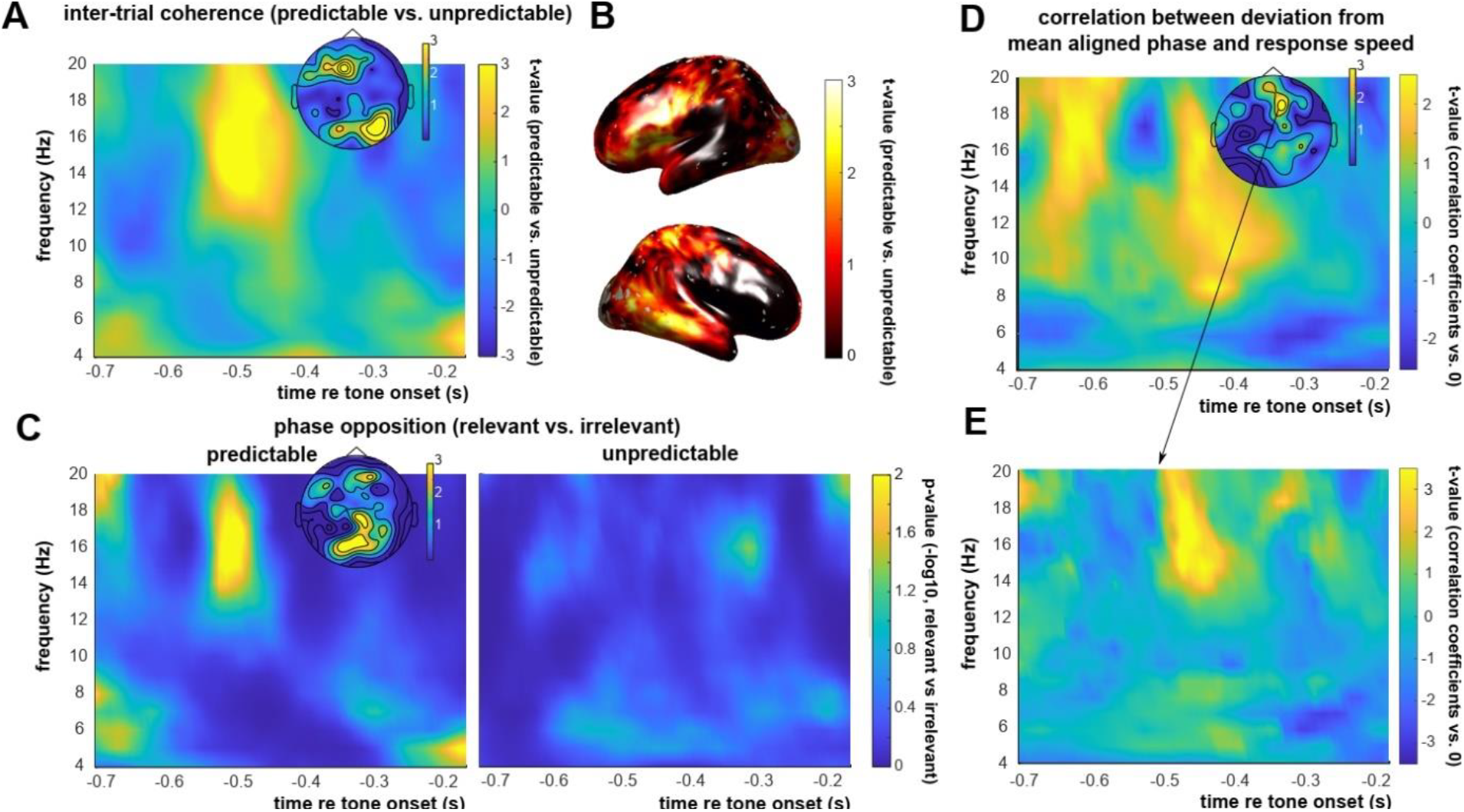
**A**. Contrast (as t-values) in inter-trial coherence (ITC) between predictable and unpredictable conditions, averaged across EEG channels. Time 0 corresponds to the onset of the second tone. The inset shows t-statistics for individual channels for -0.5s/18 Hz. **B**. Source localisation of the ITC contrast shown in the inset of panel A. **C**. Phase opposition between relevant and irrelevant conditions (−log10 of p-value), averaged across EEG channels and quantified in comparison with permuted datasets, preceding tones with predictable (left) and unpredictable onsets (right). The inset shows p-values (−log10) for individual channels for the same time-frequency point as for the inset in A. **D**,**E**. T-values from a comparison between Pearson’s correlation coefficients from individual participants and 0. Individual correlation coefficients were obtained by correlating the absolute value of the distance from the mean aligned phase with response speed in single trials. For D, correlation coefficients were averaged across EEG channels, but the inset shows t-statistics for individual channels for -0.47s/16 Hz. Results are shown for the predictable condition. E shows results for a selected EEG channel (F2, marked with an arrow in D). For visualisation, the sign of t-values in both D and E was flipped so that positive values associate deviations from the mean aligned phase with slower response speed.

### Beta oscillations align opposite phases to relevant and distracting acoustic events if their timing is predictable

If the phase alignment observed reflects a mechanism of attentional selection, then the aligned phase should be opposite, depending on whether the expected event is relevant or a distractor. We tested this hypothesis by contrasting the original data with a version permuted in a way so that its ITC is decreased if such a phase opposition between relevant and irrelevant conditions is present in the original data. In practice, this was achieved by comparing ITC, calculated separately for relevant and irrelevant conditions in the original data and then averaged, with ITC in datasets with randomly assigned condition labels to single trials (relevant or irrelevant). As the random assignment mixes the phases from relevant and irrelevant tones, opposite phases aligned to the two would lead to decreased ITC, as compared to the original data where relevant and irrelevant conditions are analysed separately. **Fig. 4C** shows the outcome from this approach.

As phase opposition can only occur in the presence of phase alignment, we selected the time-frequency points and EEG channels with an elevated ITC for predictable tones and used this selection for subsequent statistical analyses. We used a threshold of uncorrected p < 0.005 for this step, as a compromise between overly strict and liberal selection of time-frequency-channel combinations of interest. The selected combinations comprised 9 channels and included a time-frequency cluster of -0.52 – -0.44 s / 13-20 Hz.

We found reliable phase opposition between relevant and irrelevant tones that was maximal between -0.52 and -0.48 s, at frequencies between 14 and 18 Hz, and in frontal and parieto-occipital channels (e.g., O1: z = 3.55 for -0.49 s / 18 Hz; FDR-corrected p = 0.006; CPz: z = 3.73 for -0.5 s /18 Hz; FDR-corrected p = 0.006; AF4: z = 3.39 for -0.49 s /18 Hz; FDR-corrected p = 0.006). **Fig. 4C (left)** illustrates these results for the average across all EEG channels, and its inset depicts the topography for -0.5s/18 Hz. Importantly, there was no statistically reliable phase opposition (all FDR-corrected p > 0.05) prior to tones with unpredictable timing (**Fig. 4C, right**). These findings demonstrate that beta oscillations do not only align their phase to predicted acoustic events, they also adapt this phase according to the relevance of this event, just as expected from a mechanism of attentional selection.

### Single-trial deviations from phase alignment slow down pitch discrimination

In our data, the temporal predictability of a tone led to faster pitch discrimination (**Fig. 2**), a stronger alignment of oscillatory phase (**Fig. 4A**), and phase opposition between relevant and distracting tones (**Fig. 4C**). We tested whether these findings are related. We used the same selected time-frequency-channels combinations for statistical analysis.

We calculated, for each participant and various time-frequency points preceding the second tone, their mean aligned phase in relevant and irrelevant conditions, respectively. If the observed phase alignment (and associated phase opposition) is behaviourally meaningful, then a deviation from this phase in single trials should also change behaviour. For each participant, we therefore correlated the distance from the mean aligned phase with the corresponding response speed in single trials, and contrasted the obtained correlation coefficients with 0 (t-test). The outcome from this analysis is shown in **Fig. 4D** (average across all channels). For visualisation, the sign of t-values in **Fig. 4D** was flipped so that positive values associate deviations from the mean aligned phase with slower response speed. We found a reliable negative correlation between the deviation from the mean aligned phase and response speed between -0.48 s and -0.47 s and between 14 and 18 Hz, and that peaked in frontal channels (e.g., F2: t(24) = -4.35 for -0.47 s / 16 Hz, FDR-corrected p = 0.03; the inset of **Fig. 4D** shows the topographical distribution). The time-frequency representation of this neural-behavioural correlation for channel F2 (**Fig. 4E**) strongly resembled those for phase alignment (**Fig. 4A**) and phase opposition (**Fig. 4C**). Importantly, no statistically reliable correlation was observed for temporarily unpredictable tones (all FDR-corrected p > 0.05). We conclude that a deviation from the aligned phase slowed down behavioural responses, illustrating behavioural relevance of the observed putative mechanism of attentional selection.

### Visual cues do not enhance the phase alignment of beta oscillations to expected acoustic events

Finally, we tested whether the presence of a visual cue can amplify the phase alignment of neural oscillations to the expected tone timing. We first computed the ITC difference between predictable and unpredictable conditions, separately for informative and uninformative cues, and then contrasted the two with paired t-tests. We found no significant effect of the presence of an informative visual cue on the increased ITC before a predictable onset of the second tone (**Fig. S1**; all FDR-corrected p > 0.05). Informative cues did not reliably alter the phase opposition between relevant and irrelevant conditions either (all FDR-corrected p > 0.05). These findings suggest that, at least in our data, a visual cue did not contribute to the alignment of oscillatory phase to predicted tones.

## Discussion

Targets are often easier to detect and process if they occur at a predictable time^1,24^. The perceptual benefit of temporal predictability has been equally demonstrated for targets embedded in a rhythmic sequence^25,26^ and for those presented at a predictable time but without rhythmic context^27–29^, although the underlying neural mechanisms do not seem identical for rhythm- and interval-based predictions^27,28^. The link between predictive processes and neural oscillations is similarly established and takes different forms, from an anticipatory change in the oscillatory amplitude^17,30,31^ to travelling waves that are often seen as a neural correlate of predictive coding^32–34^.

For an efficient use of temporal predictability, neural resources need to be allocated to specific moments in time – to moments predicted to be most informative, or away from those predicted to be distracting. The phase of neural oscillations seems ideal for such a purpose, as it reflects the neural system’s sensitivity to sensory information^3^. An alignment of that phase according to the predicted timing and relevance of upcoming events can be a powerful mechanism of attentional selection^4^. However, experimental support for such a mechanism remained sparse, in particular in the auditory domain. In the field of “neural entrainment”, it is often assumed that neural oscillations align their phase to a rhythmic stimulus to anticipate upcoming events^6–8^. However, this assumption is difficult to verify as it is challenging to distinguish entrained oscillations from a neural response that reflects the regularity of the stimulus, rather than involving a neural process that is endogenously rhythmic^13–15,24,35^ (i.e., in the absence of a rhythmic stimulus). Moreover, in a rhythmic context, phase alignment to an event does not necessarily reflect a predictive process but can also be caused by a response to the preceding event in the stimulus sequence.

In the current study, we report evidence for a mechanism of attentional selection in time that is based on oscillatory phase. Temporal predictability did not only lead to faster reaction times during pitch discrimination, it also aligned the phase of beta oscillations to the expected stimulus, and adjusted the phase according to the stimulus’ task relevance. As the phase was aligned in a non-rhythmic context that does not bias the neural response in favour of a specific frequency, the effect is likely to involve endogenous neural oscillations. The observed phase opposition between expected targets and distractors suggests that the alignment of oscillatory phase is a neural mechanism to anticipate and select future events. Indeed, a deviation from phase alignment slowed down behavioural responses, just as expected from such a mechanism.

In paradigms that do not involve attentional competition, other studies have reported a phase alignment to predicted non-rhythmic events. Samaha et al.^36^ showed that an alignment of alpha phase to the predicted timing of non-rhythmic targets leads to more accurate perception. Daume et al.^37^ found that a similar phase alignment occurs in the delta band (∼0.5-3 Hz) during visual and tactile detection tasks. A follow-up study used transcranial brain stimulation to show that delta phase alignment causally modulates performance in such tasks^38^. A phase alignment of delta oscillations also seems to occur in anticipation of acoustic events^39^. However, in contrast to our results, two studies failed to show a phase alignment to predictable, non-rhythmic sounds during selective attention^17,20^. It is of note that both temporal predictions and competition between targets and distractors were induced cross-modally (visual vs. auditory) in these two studies. It is therefore possible that temporal predictions are strongest in the auditory system if they are induced uni-modally, in line with the finding that a visual cue did not enhance phase alignment in our study. Another possibility is that visual cues influence auditory processing (only) for typical lags between visual and auditory information in natural scenarios. For example, visual speech cues typically precede auditory ones^40^, and benefits of cross-modal input are largest if visual information precedes the auditory one by ∼30–100 ms^41^. In such scenarios, visual cues have indeed been shown to reset the phase of auditory oscillations^42,43^. It is also possible that phase-based attentional selection is restricted to situations with high competition between targets and distractors, a notion that has been raised before^17^. As competition for attentional resources seems higher within than across modalities^44^ and participants had to attend or ignore tones with similar pitch in our study, the high competition might have contributed to the successful identification of phase alignment and opposition. Finally, whereas other modalities employ the phase of alpha^16,17^ or delta oscillations^37,45^, we found that audition relies on beta oscillations. Although their paradigm did not involve a competition between targets and distractors for attentional resources, Herbst and Obleser^39^ also found a phase alignment of delta oscillations to acoustic events. Our experimental design makes it difficult to test for such slow oscillations, as their detection requires long periods without stimulus-evoked responses. However, previous work supports an important role of beta oscillations for temporal precision in auditory processing, as described in the following paragraph.

Beta oscillations are closely linked to temporal predictions, in particular in an auditory-motor network. Most of the evidence for this link comes from studies that observed an increase in beta power before or synchronised with the onset of rhythmically presented sounds^46–51^ and their omission^52^. This increase in beta power correlates with accuracy in a perceptual task^46^ and has been shown to originate from the sensorimotor cortex^22,53^, well in line with the fact that overt movements enhance precision in an auditory task^54^. Nevertheless, the phase of beta oscillations is rarely examined. Our results suggest that the observed increase in power during auditory temporal predictions goes along with an alignment of phase, where the increased power might further amplify the effect of phase on the anticipated stimulus. Although based on EEG data and therefore limited spatial accuracy, our source localisation results (**Fig. 4B**) do suggest an involvement of the right sensorimotor cortex, along with the left inferior frontal cortex that interacts with motor cortex and plays a similarly important role for auditory temporal predictions^23^. Although a method with higher spatial resolution like magnetoencephalography (MEG) is required to test this hypothesis, our data is compatible with the notion that these regions prepares the auditory system for upcoming input.

It seems a surprising finding that the alignment of beta phase occurred ∼0.5 s before the expected tone onset in our study. Although the spectral analyses required for the phase estimate necessarily introduce some temporal uncertainty, the fact that both phase alignment (**Fig. 4A**), phase opposition (**Fig. 4C**), and their correlation with behaviour (**Fig. 4D,E**) occurred at very similar times suggests that the timing of the effect is robust. In line with a possible contribution from the motor system, we speculate that the early timing could reflect a preparatory allocation of motor resources, to be available later when the auditory event needs to be encoded or suppressed (depending on its relevance). If such a preparation takes some time, this could explain why it needs to be initiated prior to the actual event. This speculative explanation is supported by the fact that temporal predictability improved the motor component (reaction time) of the pitch discrimination task, rather than its accuracy, an observation that has been made in similar form for the visual domain^55^. However, the beta phase aligned to the comparison tone that did not require a motor response. Accordingly, the phase alignment observed might neither prepare the auditory nor the motor system in isolation, but rather tune auditory-motor (including inferior frontal) networks to upcoming events, a hypothesis that can be tested, for example, in directed connectivity revealed by MEG^56^.

## Conclusion

We found that the temporal predictability of an acoustic stimulus does not only accelerate behavioural responses, it also aligns the phase of beta oscillations in the EEG to the predicted stimulus. This phase was opposite, depending on whether the stimulus is task relevant or a distractor. Deviations from the typically aligned phase slowed down pitch discrimination. These results point to a general neural mechanism to anticipate future events, where neural resources, possibly including those in auditory-motor networks, are allocated to the expected time of upcoming sensory information.

## Methods

### Participants

Twenty-six participants provided informed consent to take part in the experiment for a monetary reward of €25. The data of one participant were excluded due to a misunderstanding of task instruction. Thus, 25 participants (15 females, mean age = 27.6 years, SD = 7.8 years) were included in the final data analyses. All experimental procedures were approved by the CPP (Comité de Protection des Personnes) Ouest II Angers (protocol number 2021-A00131-40).

### Stimuli and Experimental Design

Participants performed a pitch discrimination task in which three tones were presented consecutively in each trial (**Fig. 1**). They were instructed to compare the final tone (reference tone) with either the first or second tone (comparison tones) in each trial. The identity of the comparison tone (first or second tone) was kept constant within an experimental block. Each trial was ∼5.5 s – 6.5 s long (including a response time window of 2 s) and there was a randomly selected interval between 0.7 s and 2 s between trials. Each block consisted of 36 trials (average block duration: 4.62 minutes) and participants completed ten blocks.

The pitch of the first tone was randomly selected between 392 Hz (G4 musical note) and 783.99 Hz (G5) to prevent forming a memory trace of a specific pitch throughout the experiment. Pitch differences between the three tones were expressed in terms of “units”. One unit corresponds to the pitch discrimination threshold that was determined separately for each participant and for the pitch difference between the reference tone and each of the two comparison tones (first and second tones) in an adaptive staircase procedure before the main experiment (see next section). In each trial, the pitch of the comparison tone differed from that of the reference tone by +2, +1, 0, -1, or -2 units. The pitch of the second tone differed from that of the first tone +1, 0, or -1 units, where the unit refers to the one between reference and comparison tones in the same trial. All pitch differences were varied across trials and selected pseudo-randomly. The stimuli were presented at a comfortable listening level (ca. 70 dB SPL).

Our paradigm (**Fig. 1**) included the experimental manipulations of temporal predictability (varied by block), task relevance (varied by block), and cue informativeness (varied by trial).

Temporal predictability was operationalized as the interval between the first and second tones. In some blocks, the second tone always occurred 1.5 s after the first tone (predictable condition). In other blocks, the interval was randomly chosen from a range between 1 and 2 s (unpredictable condition). The third tone always occurred 1.5 s after the second tone.

The task relevance of the second tone was manipulated by varying the identity of the comparison tone. When the second tone was the comparison tone, it was relevant. When the first tone was the comparison tone, the second tone was an irrelevant distractor. Participants were informed of the identity of the comparison tone before the start of each block.

Cue informativeness was varied in the form of a visual cue. This cue appeared 1 s before the onset of the first tone, and disappeared with its offset. The cue could be either informative or uninformative for the pitch difference between the first and second tones. Cues were not informative for the pitch difference between the first two tones and the reference tone. The informative cue (**Fig. 1**) was an arrow that pointed either to the upper right corner (second tone higher in pitch), right side (second tone same in pitch), or the lower right corner (second tone lower in pitch). For the three uninformative cues, the positions of the short lines of the arrows were randomly placed such that there was no directional element in the symbol anymore. The three uninformative cues were presented with the same probability for each pitch difference between first and second tones.

The participants pressed ‘F’ or ‘J’ on the keyboard for ‘higher’ or ‘lower’ responses (counterbalanced across participants), and the spacebar for a ‘same’ response. The response time window was 2 s long, and the next trial started automatically.

The stimuli were presented via Matlab 2019a (MathWorks, Inc., Natick, USA) and Psychtoolbox^57^. Auditory stimuli were presented with a Fireface UCX soundcard and Etyomic ER-2 inserted earphones. Synchronization between sound and EEG data was ascertained by using the same soundcard for triggers sent to the EEG amplifier.

### Adaptive Staircase Procedure

Individual pitch differences (i.e. “units”) between reference and comparison tones were determined in a 1-up-1-down adaptive staircase procedure using the Palamedes toolbox^58^. These pitch differences were determined separately between the reference tone and each of the two comparison tones (first and second tones) to account for the different time intervals between them (first tone to reference tone: 2.5-3.5 s; second tone to reference tone: 1.5 s). The order of adapted pitch difference (first or second tone vs. reference tone) was counterbalanced across participants. The second tone was not presented for the adaptation of the pitch difference between first and third tone (i.e. the time interval between tones was identical to the main experiment, but there was no distractor). The pitch of the comparison tone was fixed at the mean of the pitch frequency range used in the main experiment (i.e., 588 Hz).

In each trial of the staircase procedure, participants had to indicate whether the reference tone was higher than, lower than, or identical in pitch as the comparison tone. The pitch difference between the two decreased if participants had correctly indicated the direction of a pitch difference (higher or lower), and increased if participants had responded with the incorrect direction (e.g., “lower” response when the reference tone was higher in pitch). Trials with identical pitches were discarded. The adaptive procedure stopped after 9 reversals and the last 6 reversals were used to calculate individual pitch difference thresholds.

### EEG Recording and Data Processing

The Biosemi Active 2 amplifier (Biosemi, Amsterdam, Netherlands) with 64 active electrodes positioned according to the international 10-10 system was used for EEG recording, with a sampling rate of 2048 Hz. Instead of more typical reference and ground electrodes, common mode rejection was achieved by the “Common Mode Sense” active electrode and a “Driven Right Leg” passive electrode. Signal offsets for all electrodes were kept below 50 μV.

All EEG pre-processing steps were conducted using the fieldtrip toolbox^59^ in Matlab 2021a (MathWorks, Inc., Natick, USA). EEG data were re-referenced to the average of all EEG electrodes and high-pass and low-pass filtered with cut-off frequencies of 1 and 40 Hz, respectively (4th order Butterworth filter). An independent component analysis (ICA) was applied to down-sampled data (sampling rate: 256 Hz) to identify artefacts such as eye blinks and muscle artefacts. Noisy channels and contaminated ICA components were identified with visual inspection and removed from data at the original sampling rate.

The continuous EEG data were segmented into trials between -2 s and +5 s relative to the onset of the first tone. Segments with an absolute amplitude exceeding 100 μV were rejected. As all of our experimental manipulations were designed to change neural processing of the second tone, we re-epoched data from -1.5 s to +1.5 s relative to the onset of the second tone.

For these epochs, we used Fast Fourier Transformation (FFT) to estimate single-trial phases for time points from -0.7 s to 0 s (relative to the onset of the second tone; steps of 0.01 s) and frequencies from 2 Hz to 20 Hz (steps of 1 Hz). Each time point refers to the centre of a time window subjected to FFT, with the linearly spaced window sizes ranging from 2 cycles (for 2 Hz) to 4.4 cycles (for 20 Hz) for each frequency.

### Data and Statistical Analyses

We used the estimated phases to quantify the hypothesized phase alignment at a range of time-frequency points preceding the second tone. Phase alignment was quantified using inter-trial coherence (ITC):

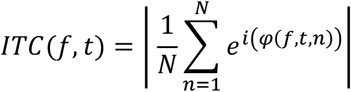

where *φ(f,t,n)* is the phase in trial *n* at frequency *f* and time point *t*, and *N* is the number of trials. For a given time-frequency point (*t,f*), ITC is high if its phase is consistent across trials, and ranges between 0 (no phase consistency) and 1 (perfect phase consistency).

To test whether the phase of neural oscillations aligns to temporarily predictable sounds, we used a paired t-test to contrast ITC between predictable and unpredictable conditions (**Fig. 4A**). We expected higher ITC for predictable sounds. The hypothesized phase opposition between relevant and irrelevant conditions would lead to reduced ITC (as compared to phase alignment without phase opposition) if trials are pooled across these two conditions before ITC is computed. Such an effect is of particular relevance in the presence of strong phase alignment in each of the conditions, as opposite phases are combined and lead to phase cancellation. For both predictable and unpredictable conditions, ITC was therefore computed separately for relevant and irrelevant conditions and then averaged across relevance, avoiding phase cancellation.

To test for phase opposition between relevant and irrelevant conditions, we leveraged the fact that such phase opposition reduces ITC (as compared to phase alignment without phase opposition) if trials are pooled across conditions, but this reduction does not occur if ITC is computed separately for each condition and then averaged. Separately for predictable and unpredictable conditions, we again computed ITC for relevant and irrelevant conditions and averaged across relevance. We then compared the outcome with a version where the relevance label (relevant or irrelevant) is randomly assigned to trials. Only in the presence of phase opposition, the random assignment would reduce ITC (as compared to the original data) as trials with opposite phases are pooled. The random assignment (and ITC computation) was done 1000 times to obtain a distribution of ITC values that would be obtained in the absence of phase opposition. For each time-frequency point, the original data was then compared with the permuted data to obtain group-level z-scores:

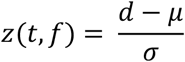

where *z* reflects the group level effect in the original data, *d* is the ITC in the original data at time *t* and frequency *f*, and *μ* and *σ* are mean and standard deviation (across permutations) of the subject-averaged ITC at the same time-frequency point in the permuted datasets, respectively. Z-scores were then converted to p-values and are shown in **Fig. 4C** (e.g., z = 1.645 corresponds to a significance threshold of α = 0.05, one-tailed due to the directional hypothesis). We expected higher ITC in the original data as compared to the permuted datasets, but only for temporarily predictable tones.

To test whether the visual cue helped participants align the phase of oscillations to expected sounds, we computed the ITC difference between predictable and unpredictable conditions and contrasted this difference between informative and uninformative cues with paired t-tests (**Fig. S1**). We expected a larger ITC difference after informative cues. We also contrasted our measure of phase opposition between informative and uninformative cues. This contrast required z-scores (original vs. permuted data) on the individual (rather than group) level which we obtained by applying the analysis described above to individual participants. Z-scores were then contrasted between informative and uninformative cues using a paired t-test. We expected higher z-scores for informative cues.

If phase alignment contributed to pitch discrimination performance, then single-trial deviations from the typically aligned (i.e. mean) phase should lead to a change in performance. We focused on response speed, as temporal predictability led to faster but not more accurate responses. We calculated, for each trial, participant, and time-frequency point, the circular distance to the mean aligned phase in the respective condition (predictable-relevant, predictable-irrelevant, unpredictable-relevant, unpredictable-irrelevant). We then correlated the absolute value of these single-trial phase differences with single-trial response speed, yielding one correlation coefficient for each participant and time-frequency point. For this correlation, trials from relevant and irrelevant conditions were pooled, as a deviation from the respective mean phase should change response speed irrespectively of relevance. Predictable and unpredictable conditions were analysed separately, as we expected to find a correlation only in the presence of phase alignment (and hence only for predictable tones). We used t-tests to compare these correlation coefficients with 0, to draw conclusions on the group level. We expected a negative correlation between deviation from the mean aligned phase and response speed and implemented this hypothesis in the form of one-tailed tests.

Paired t-tests were used to contrast neural responses at electrode Fz (an electrode that typically shows strong activation in auditory experiments, as illustrated in the topographies of **Fig. 3**), prior to and evoked by the second tone, between conditions.

All multiple comparisons were corrected using the false discovery rate (FDR).

We designed a linear mixed model (**Fig. 2**) to quantify the effects of temporal predictability, task relevance and cue informativeness on behavioural performance (reaction time and accuracy), with participant included in the model as the random intercept. For all analyses (including the correlation with neural data), we converted reaction time to response speech (its reciprocal) to increase the intuitiveness of results (higher values = faster responses). We verified that our results did not depend on this choice.

Finally, we explored the neural origins of the phase alignment found in the EEG data. For this purpose, we used the fieldtrip toolbox to construct a standard volume conduction model and combined it with EEG electrode locations to calculate the leadfield matrix (10 mm resolution). We calculated a spatial common filter using the LCMV beamformer method^60^ (lamba = 5%), using data from -1.5 s to +1.5 s relative to the onset of the second tone (all conditions pooled). This resulted in 2,015 source locations that were inside the brain.

The spatial common filter was then applied to Fourier-transformed single-trial data (output from the FFT described above) from individual participants. The spatially filtered, Fourier-transformed single trials were combined to form ITC, using the formula provided above. This step yielded one ITC value for each of 2,015 voxels inside the brain per participant. The result of this analysis is shown in **Fig. 4B**. Note however that, due to the low spatial resolution of EEG, we consider our source-level results as explorative and not necessarily statistically informative. The statistical analysis of our main hypotheses is done on the sensor level (described above).

## Acknowledgements

This work was supported by a grant from the Agence Nationale de la Recherche (ANR-21-CE37-0002 to B.Z.).

## Supplemental Figures

**Figure S1.**
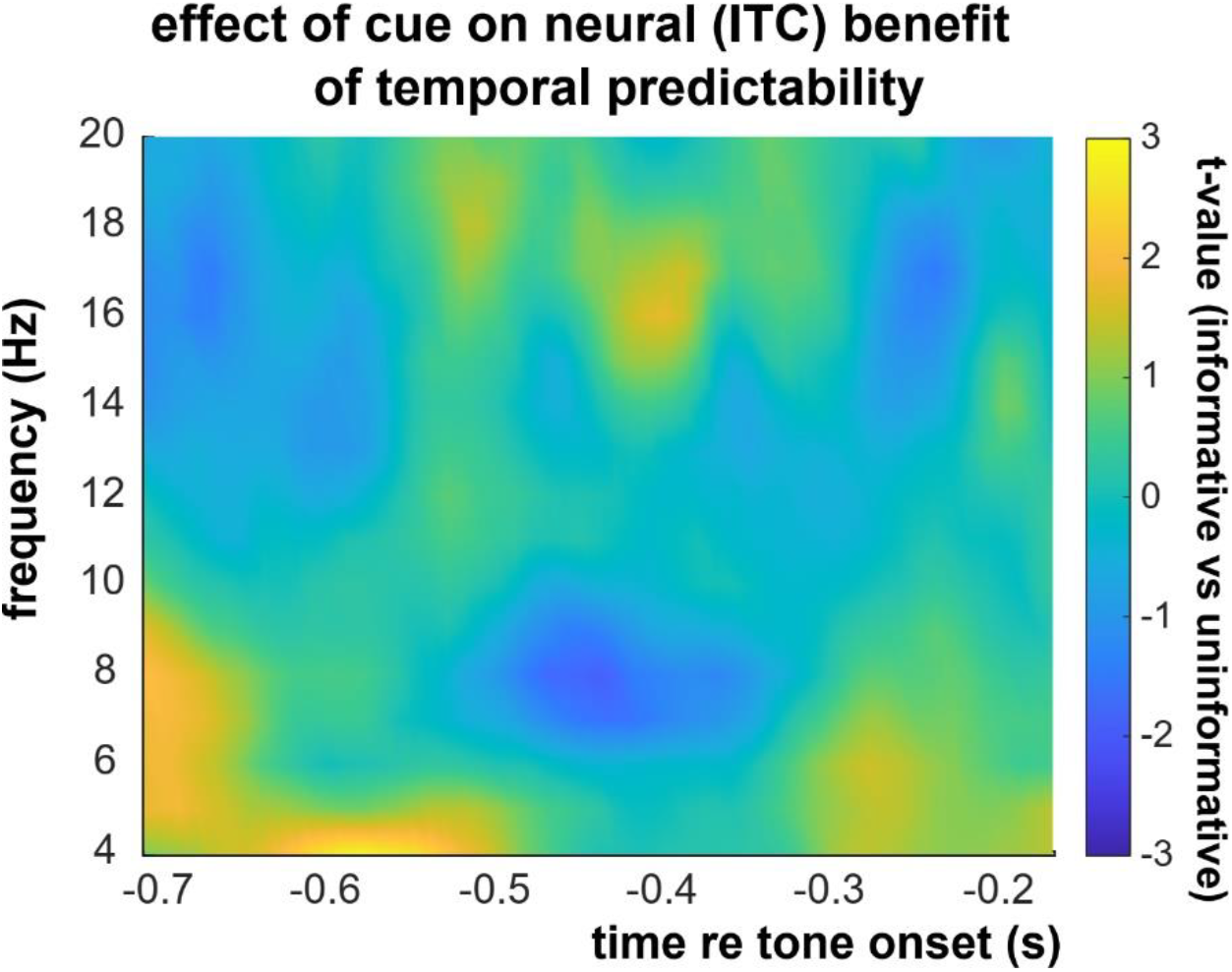
Contrast (as t-values) in the ITC difference between predictable and unpredictable conditions (averaged across EEG channels), between informative and uninformative visual cues.

